# Plasma untargeted metabolomics reveals distinct metabolic disturbances between sexes following inhalation of ultrafine particles and exposure to aircraft noise in mice

**DOI:** 10.64898/2026.10.27.762145

**Authors:** Thuy Thi Lai, Aino-Kaisa Piironen, Elena Puris, Veronika Górová, Mika Ihalainen, Tuukka Kokkola, Jiayin Zheng, Thomas Münzel, Andreas Daiber, Pasi Jalava, Olli Sippula, Katja M. Kanninen

## Abstract

Traffic noise and air pollution, particularly ultrafine particles (UFP), are significantly associated with health risks. However, the systemic metabolic consequences of the combined effects of UFP and aircraft noise (AN) exposure on health remain poorly understood. We aimed to investigate changes in plasma metabolites in healthy wildtype C57BL/6J mice following whole-body inhalation exposure to UFP (average diameter <40 nm), AN and their combination (UFP+AN). Liquid chromatography-mass spectrometry (LC-MS)-based untargeted plasma metabolomics analysis was undertaken for UFP, AN, and combined UFP+AN-exposed mice of both sexes. This study identified a total of 1146 plasma metabolites. Across exposure groups, AN induced the greatest plasma metabolomic changes in both sexes, while UFP significantly altered metabolites only in females, indicating a sex-specific response. The combined UFP+AN exposure produced a modestly enhanced effect in males compared with either single exposure but showed an attenuated effect in females. Pathway analyses revealed that lipid and amino acid metabolism were the most affected, followed by xenobiotics, and cofactors and vitamins, after exposure in both female and male mice. Additionally, in the clean air control group, we identified sex differences in 49 plasma metabolites, which mostly included lipids and amino acids. Following exposure, however, sex-dependent metabolic differences became more pronounced and exposure-specific, suggesting that UFP and AN have distinct effects on plasma metabolome of male and female mice that extend beyond baseline sexual dimorphism observed in unexposed animals. To our knowledge, this study is the first comprehensive characterization of systemic metabolic changes in mice after combination of UFP and AN exposure, providing new insight into the interplay between traffic noise and air pollution health risks.

## Introduction

Health impacts and disease burden relating to environmental air and noise pollution exposure have been increasingly researched over the last decade. Particulate matter (PM) air pollution and aircraft noise (AN) have been identified as two major environmental stressors and risk factors for several disorders ^1–4^. According to the World Health Organization (WHO), air pollution contributed to 6.6 million deaths worldwide in 2021, with the largest impacts seen in the low- and middle-income countries where people often have higher exposure risks and limited access to health care, while noise exposure is also estimated to result in the loss of over one million healthy life years in Europe ^5,6^. The simultaneous presence of air pollution and AN is a significant concern in real-world environments. Furthermore, people living close to busy roads, industrial areas, airports, or under flight paths may have a higher risk of exposure to the co-occurrence of both air pollution and AN, which negatively impacts their quality of life and various adverse health issues ^7^.

Epidemiological studies have demonstrated the association of air pollutants or traffic noise to adverse health effects ^4,8–11^. Exposure to inhaled soot ultrafine particles (UFP, particle diameter less than 100 nm) is associated with an increase in mortality and hospital admissions ^12^. Chronic exposure to average noise levels exceeding 55 decibels (dB) is also linked to adverse health outcomes, including hypertension, ischemic heart disease, and significant sleep disturbances, and may exacerbate cardiovascular disease and psychological stress ^11^. Exposure to air or noise pollution is also linked to dementia and cognitive impairment ^13,14^. Furthermore, a recent study reported that chronic exposure to intense AN among military personnel was associated with noise-induced hearing loss, increased oxidative stress, inflammation, and impaired cognitive performance ^15^.

Both *in vivo* and *in vitro* studies have provided insights into the mechanisms underlying harmful exposure effects caused by either air or noise pollution ^10,16–20^. However, the underlying molecular and cellular mechanisms that mediate co-exposure effects of air pollution particles and noise remain unclear. A paper by Kuntic *et al.* described the combined effects of exposure to urban PM_2.5_ and AN to cause additive damage reflected by endothelial dysfunction, increased blood pressure, oxidative stress, and inflammation in mice with differential effects in lung, brain, aorta and heart tissues ^21^. In addition, a recent paper by Ameli *et al.* provided a deeper analysis of the proteomic changes in animal tissues from the Kuntic *et al.* study ^22^. This analysis revealed that the combined exposure triggered the strongest immune and oxidative stress response in the lungs, while the effects on brain and heart were significantly smaller ^22^.

One emerging technology to capture the biological response to environmental exposures is plasma metabolomics. It is known that metabolic changes directly reflect the response to external exposures and disease progression ^23^, and therefore, measuring metabolic changes after exposure has the potential to reveal toxicity and other underlying cellular and molecular mechanisms following exposures. A previous study has reported significant changes in the plasma metabolome, mostly related to phospholipids, in individuals exposed to PM_2.5_ pollution over one-year follow-up period ^24^. This finding aligns with an earlier study showing that short-term PM_2.5_ exposure disrupts various plasma metabolic pathways in young adults in China ^25^. In addition, exposure to UFP also resulted in alterations to the plasma metabolome of people living in Boston, USA ^26^. Similarly, exposure to noise also led to alterations in the plasma metabolites ^27,28^. However, insight into the effects of combined UFP and noise exposure on the plasma metabolome remains limited.

Another knowledge gap remains in understanding sex-specific responses to environmental stressors ^16^. Most of the existing UFP and/or AN exposure literature has relied on male or female-only cohorts, and only one study has reported sex-specific responses in obese mice after exposure to UFP ^29^. The authors demonstrated that obese female mice may be at higher risk of metabolic dysfunction during exposure, in comparison to males. To date, no study has examined whether combined UFP and AN exposure produces sex-associated differences in systemic metabolism. Given that sex differences in oxidative stress, lipid metabolism, and cardiovascular or neurodegenerative disease risk are well documented ^30–32^, this aspect represents an important evidence gap in research of environmental health.

This study addresses the above-mentioned knowledge gaps by using untargeted metabolomics to characterize plasma metabolite profiles observed under three different exposure conditions in healthy mice: UFP alone, AN alone, and combination of UFP and AN exposure. We further examined whether UFP and/or AN exposure induces sex-specific metabolic changes by identifying metabolic pathways across exposure conditions and sexes. This study aims to clarify the systemic biological consequences of a real-world co-exposure scenario and provide insightful knowledge into health vulnerabilities in the context of the interplay between traffic noise and ultrafine air pollution.

## Materials and Methods

### 1. Mouse husbandry

Wildtype 3-month-old female (n=40) and male (n=24) littermates with C57BL/6J (The Jackson Laboratory) background were used for the experiments. Mice were kept in a controlled environment with a 12-hour light/dark cycle, consistent humidity and temperature and ad libitum access to food and water. All animals were group-housed (up to 6 mice per cage), with male and female mice housed separately. Within each sex, animals were evenly distributed across conditions. All animals were cared for by the laboratory animal center at the University of Eastern Finland (animal license number ESAVI-2024-032320).

### 2. Exposure

A whole-body inhalation chamber utilizing the mini combustion aerosol standard (miniCAST) burner ^33,34^ running on propane was used to expose mice to UFP. Briefly, a propane (mixed with nitrogen)/air co-flow laminar diffusion flame was quenched by nitrogen and the resulting particles were transported into a nitrogen stream, then travelled into a whole-body animal exposure chamber designed to simulate real-world conditions (Supplementary Fig. 1A). We used a playback of recorded aircraft events from German Airports, Germany ^35^ to simulate real-life exposure conditions to aircraft noise. The mice were randomly divided into four groups: clean air (control), freshly generated combustion-derived ultrafine particles (UFP), aircraft noise (AN), and a combination of UFP and AN (UFP+AN). The exposures took place on weekdays for 4 hours each day, for a total of 9 days over the course of 2 weeks. Mice were weighed prior to exposure, and then again one week after exposure and on the day following the final exposure (Day 10). The geometric mean size of the CAST-burner generated UFP was 32.63 ± 1.29 nm, with an average total concentration of 8.37 × 10□± 2.59 × 10 particles/cm³ per day. The mass concentration was 73.18 ± 7.41 µg/m³, and approximately one-fifth of the total mass was black carbon (14.53 ± 2.58 µg/m^3^). The aircraft noise exposure featured a 2-hour long noise pattern of 69 aircraft noise events with a duration of 43 seconds each and a maximum sound pressure level of 85 dB(A) and an average sound pressure level of 72 dB(A), as previously described ^35^.

### 3. Transmission electron microscopy (TEM)

TEM samples were obtained from the stabilizing chamber connected to the CAST burner using an aspiration sampler with 0.3 Lpm flow rate, following the method described by Leskinen *et al.* ^34^. PM samples were collected on holey carbon grids (S147-4 Holey carbon film 400 Mesh Cu; Agar Scientific Inc., USA), and the morphology of these particles was examined using TEM (JEM-2100F, JEOL Inc., Tokyo, Japan).

### 4. Mouse blood collection and plasma isolation

On the day following the final exposure day (Day 10), when the mice were 3.5 months old, mice were first anaesthetized by intraperitoneal injection of 25 mg/mL of Euthbard vet. Blood was then collected by intracardiac puncture in the right ventricle into a 1 mL syringe containing 25 mM EDTA (Invitrogen, cat#15575020) and immediately placed on ice. Blood was then centrifuged at 4°C for 10 min at 2000 × g to enable separation of plasma from blood cells and platelets. After that, the supernatant containing plasma was collected and further centrifuged at 4°C for 10 min at 2000 × g to completely deplete platelets in the plasma sample. Plasma was transferred into new Eppendorf tubes and aliquoted at 150 µl per tube to barcoded tubes provided by Metabolon Inc and then stored at -70°C until use.

### 5. Plasma sample preparation and Ultrahigh Performance Liquid Chromatography-Tandem Mass Spectrometry analysis

The plasma sample preparation was performed at the Metabolon platform as described previously ^36–39^. Briefly, samples were prepared using the automated MicroLab STAR® system from Hamilton Company. Several recovery standards were added prior to the first step in the extraction process for QC purposes. To remove protein, dissociate small molecules bound to protein or trapped in the precipitated protein matrix, and to recover chemically diverse metabolites, proteins were precipitated with methanol under vigorous shaking for 2 min (Glen Mills GenoGrinder 2000) followed by centrifugation. The resulting extract was divided into multiple fractions: two for analysis by two separate reverse phase (RP)/UPLC-MS/MS methods with positive ion mode electrospray ionization (ESI), one for analysis by RP/UPLC-MS/MS with negative ion mode ESI, one for analysis by HILIC/UPLC-MS/MS with negative ion mode ESI, while the remaining fractions were reserved for backup. Samples were placed briefly on a TurboVap® (Zymark) to remove the organic solvent. The sample extracts were stored overnight under nitrogen before preparation for ultra-high-performance liquid chromatography-tandem mass spectrometry (UHPLC-MS/MS) analysis in both positive and negative ionization modes. The MS analysis alternated between MS and data-dependent MS^n^ scans using dynamic exclusion. The scan range varied slightly between methods but covered 70-1000 m/z.

### 6. Compound identification and data pre-processing

Raw data were extracted, peak-identified, QC processed, quantified, and annotated by the Metabolon pipeline (Integrated Bioinformatics Platform, Version 3.0, Metabolon). Metabolites were identified by comparison to a proprietary library of over 5,400 purified standards based on retention time/index, mass-to-charge ratio (m/z), and MS/MS spectral data. An additional 7000 mass spectral entries have been created for structurally unnamed biochemicals, which have been identified by virtue of their recurrent nature (both chromatographic and mass spectral). Next, the data were batch-and volume-normalised to median scale, and missing values were imputed using the minimum value across all batches in the median-scaled data. Lastly, the normalized and imputed data were log-transformed for statistical analysis.

### 7. Bioinformatics and Statistical Analysis

Statistical analysis was performed in R (v. 4.4.2; R Core Team, 2024). Preprocessed data were analysed using linear regression modelling with the *limma* package v.3.62.2 ^40^. All samples were included in the model, which was adjusted for the animal weight after exposure, as plasma metabolite levels are known to be influenced by total body weight ^41^. We therefore conducted analysis by adjusting metabolite levels to body weight at the time of blood collection (Day 10 post exposure). Differences in metabolites between each exposure group and the control group were examined using the makeContrasts() and contrasts.fit() functions in addition to the basic lmFit() and eBayes() functions in *limma*. Additional models were run to identify sex differences and were adjusted for the post-exposure weight. Furthermore, statistical analyses used models appropriate for unbalanced sample sizes. Contrasts were then applied to male vs. female samples for each experimental group (control, UFP, AN, UFP+AN). In all regression models, multiple testing correction for p-values was conducted using Benjamini-Hochberg’s method ^42^ to control False Discovery Rate (FDR), and the results with an adjusted p-value (adj. p) <0.05 were considered statistically significant. Weight differences between pre- and post-exposure measures and sexes were examined using two-way repeated measures ANOVA followed by Tukey’s post hoc test. Data were visualized using principal component analysis and the R packages *ggplot2* (v.4.0.3), *ComplexHeatmap* v.2.22.0 ^43^, and *circlize* v.0.4.16 ^44^.

## Results

### Distinct metabolic response signatures of UFP, AN and combined exposure compared to controls

Following 9 days of exposure to freshly generated UFP, AN, and UFP+AN over a 2-week period, mice were sacrificed at day 10 for blood collection and plasma metabolomic analysis (Fig. 1A). Mouse body weights were unaffected by UFP and/or AN exposure in both female and male mice at one week of exposure and the day 10 endpoint (Supplementary Fig.1E-G), indicating that the exposure setup did not cause gross toxicity. TEM micrographs confirmed the shape and size of the CAST burner generated particles (Supplementary Fig. 1B) and the size distribution, and AN exposure pattern were verified as shown in Supplementary Fig. 1C, D.

**Fig. 1:**
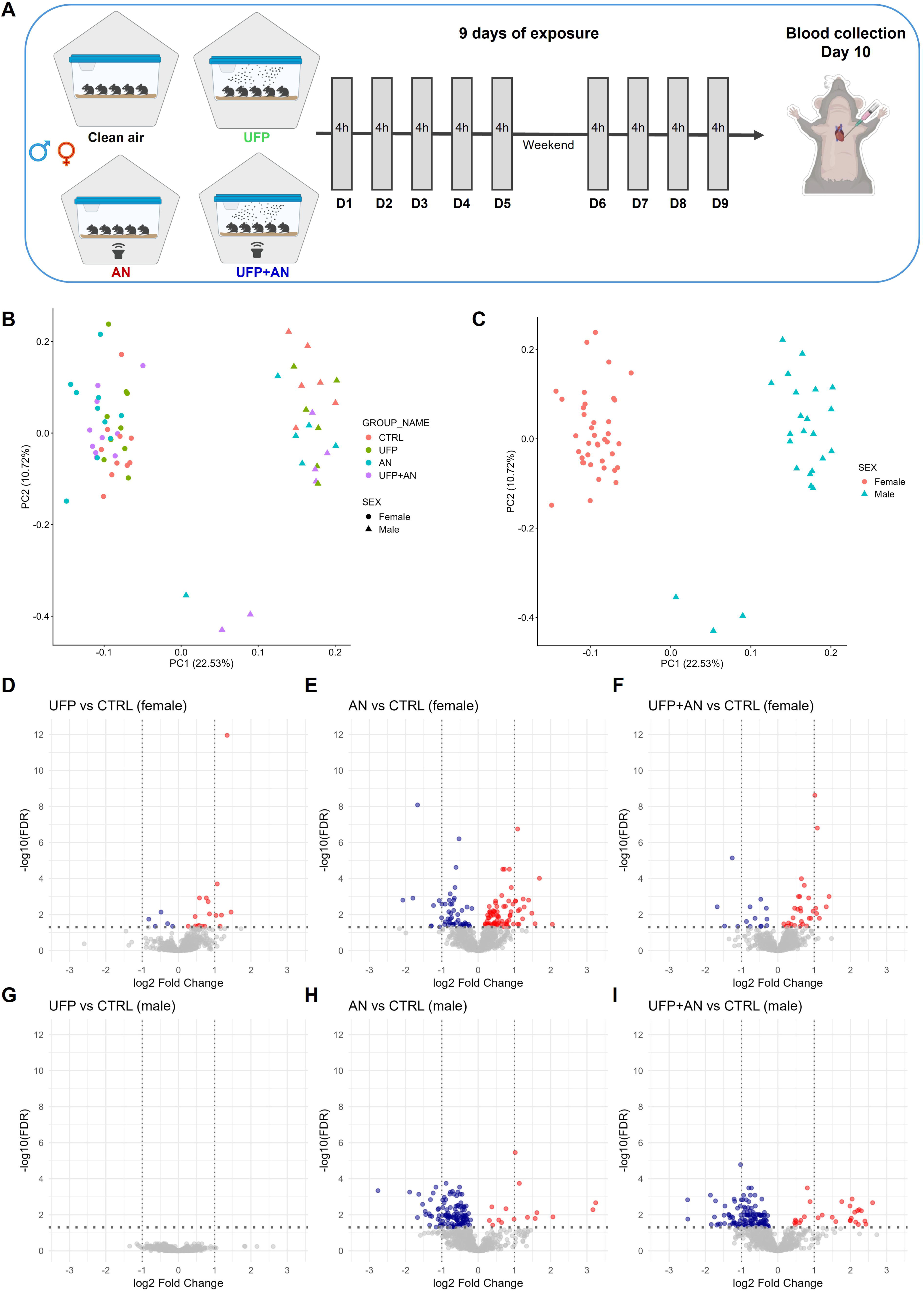
Plasma metabolite alterations in ultrafine particle (UFP), aircraft noise (AN) and co-exposed (UFP+AN) mice compared to controls by sex. A) Schematic illustration of animal exposure. The metabolite profile overview of B) the experimental groups, and C) female (red) and male (blue) mice in principal component analysis. D-I) Volcano plots for significantly (adj. p < 0.05) altered n=10 females per group and n= 6 males per group.

A total of 1146 biochemicals were detected in plasma samples across exposure groups and in both sexes (Supplementary Table 1). Among these, 1024 metabolites were annotated, while 122 were unnamed. The overall differences and similarities of the plasma metabolome across exposure conditions and sexes were assessed using the unsupervised multivariate statistical method, principal component analysis (PCA). The PCA plots displayed a clear distinguishable metabolic profile in females compared to males across the exposure conditions (Fig. 1B, C).

We first assessed statistically significant metabolic alterations specific to each exposure group in comparison to the control group in both female and male mice (adj.p <0.05). Exposure to UFP resulted in significant alterations of 22 plasma metabolites in females, of which 5 metabolites were decreased and 17 increased compared to controls (Fig. 1D and Fig. 2A). In contrast, no significant changes in metabolites were observed in the plasma of males after UFP exposure (Fig. 1G and Fig. 2A). Exposure to AN led to distinct metabolic responses between the sexes (Figure 1E, H). In females exposed to AN, a total of 135 metabolites were significantly altered, out of which 81 were increased and 54 reduced (Fig. 1E). Among the altered metabolites, 37 were classified as amino acids, and 30 belonged to a lipid pathway. The remaining metabolites included peptides, nucleotides, xenobiotics, cofactors and vitamins, and energy/citric acid (TCA) cycle pathways (Fig. 2B). Conversely, a greater number of metabolites were decreased (124 metabolites) compared to those that were increased (15 metabolites) in males exposed to AN, resulting in 139 altered metabolites overall (Fig. 1H). They included 76 lipids, 23 amino acids, 16 xenobiotics, 7 cofactors and vitamins, and 2 nucleotides (Fig. 2B). When concomitantly exposed to both UFP and AN, male mice exhibited 156 significantly altered plasma metabolites— 128 decreased and 28 increased, including 85 metabolites from lipids, 23 xenobiotics, 22 amino acids, and 6 cofactors and vitamins (Fig. 1I and Fig. 2C). In comparison, females showed changes only in 45 plasma metabolites, including 11 amino acids, and a smaller number in lipid, xenobiotics, cofactors and vitamins under the same combined exposure (Fig. 1F and 2C). These results suggest that AN exposure caused the largest number of metabolomic changes in both sexes, while the changes after UFP exposure were more pronounced in females in comparison to males.

**Fig. 2:**
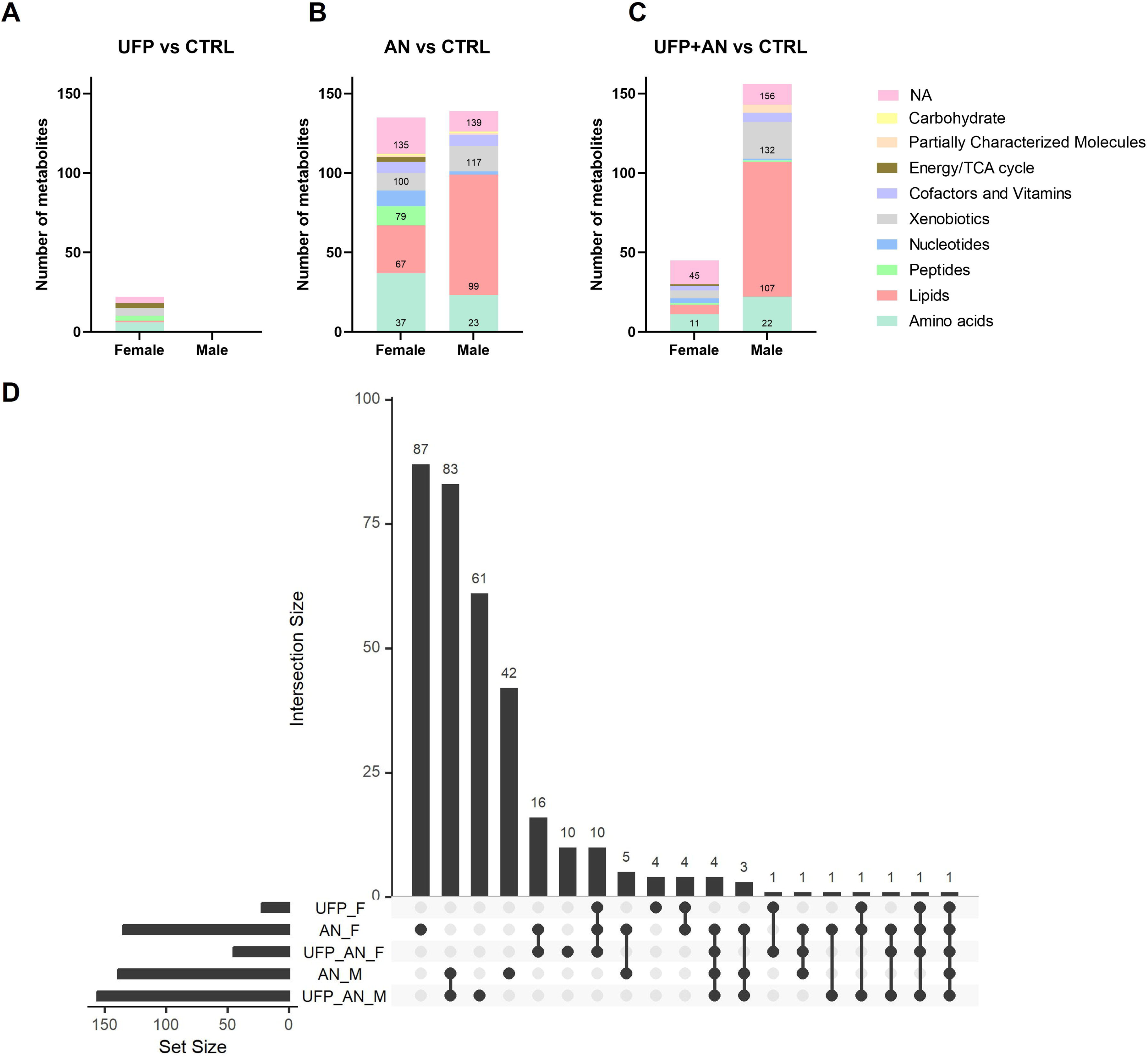
Differential metabolite profiles following UFP, AN and UFP+AN exposures. A-C) The number of significantly altered metabolite in each super pathway involved following UFP, AN and UFP+AN exposures. D) Upset plot for similarly altered metabolites across exposure groups. n=10 females per group and n=6 males per group.

Using the significantly altered metabolites following the exposures, we explored shared and unique metabolites across exposure conditions and sexes (Fig. 2D). Overall, we identified highly distinct metabolite changes between the sexes in each exposure condition. In females, we observed 87 significant unique biochemicals after AN exposure, 4 unique metabolites (phosphoethanolamine, argininosuccinate, aspartylisoleucine, and unknown X-25265) from UFP exposure, and 10 (e.g., creatinine, glycerol 3-phosphate, and inosine) from UFP+AN exposure (Fig. 2D). Additionally, 12 metabolites (e.g., amino acids imidazole lactate, 5-hydroxylysine, pro-hydroxy-pro) were altered in all three exposure conditions, of which 10 were significantly changed only in females. Furthermore, females showed an additional 21 common metabolites shared across two exposure conditions; 16 in AN and UFP+AN exposure, 4 in UFP and AN, and 1 (N-acetylalanine) in UFP and UFP+AN exposure, which were not altered in males. Considering that no significant metabolite alterations were observed in UFP exposure condition in males, shared and unique metabolites were assessed for only AN and UFP+AN exposures. There were 42 significant unique metabolites detected in the AN exposure group, 61 unique metabolites in the UFP+AN exposure, and 83 common biochemicals in both groups, which were only altered in males (Fig. 2D).

Together, these results indicated that the combination of UFP+AN exposure caused the largest number (156 metabolites) of metabolomic changes in WT males, suggesting a modest enhancement effect of the combined exposure. Additionally, AN exposure was the main trigger to plasma metabolome change in both sexes.

### Sex-dependent differentially abundant metabolites across UFP, AN and combined exposure

As we observed the distinct metabolic profile across all experimental groups and sexes, we next directly compared metabolites between males and females in the corresponding exposure conditions. Further analysis and comparison revealed slight differences in the plasma metabolome between unexposed female and male C57BL/6J mice (Fig. 3). A total of 49 metabolites were found to differ by sex, accounting for 4.3% of the total 1146 detected metabolites. Specifically, 30 metabolites were present at higher levels in the 3-month-old males than in females of the same age at baseline, whereas 19 metabolites were present at lower levels in males than in females (Fig. 3A). Among these, 21 metabolites belonged to the lipid pathway and 8 to the amino acid pathway, while the remaining metabolites were assigned to other metabolite groups or remained unannotated (Fig. 3E).

**Fig. 3.**
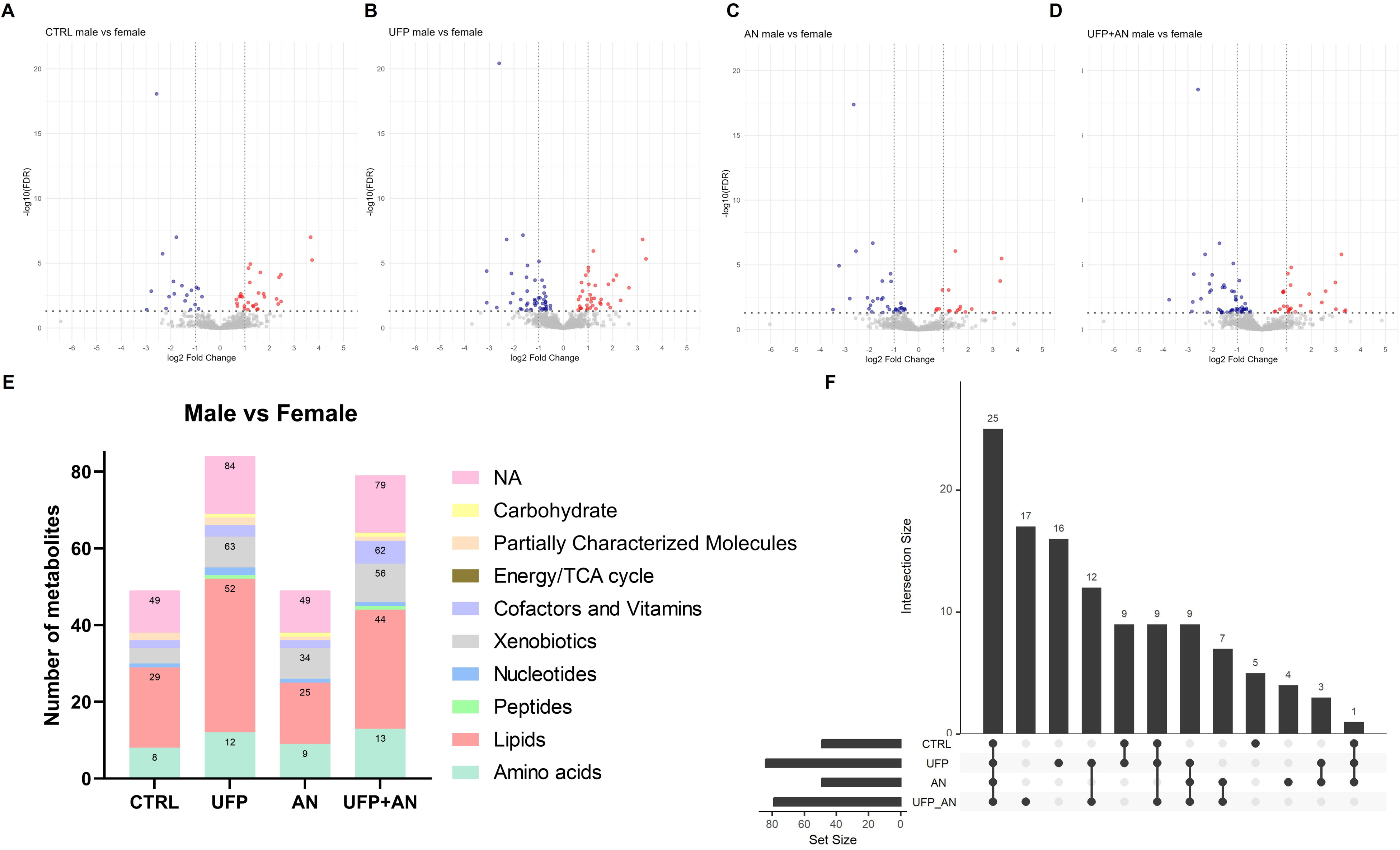
Comparisons of differentially abundant metabolites determined in male vs female mice. A-D) Volcano plots for sex-dependent metabolic response. E) The top altered super pathways involved in following exposures in males vs females. F) Upset plot comparing overlaps of altered metabolites between sexes.

Despite the small baseline sex difference in plasma metabolome, exposure to UFP alone and with AN was associated with a marked increase in metabolites showing sex differences (Fig. 3B-E). Most significantly altered metabolites were classified as lipids, amino acids, and xenobiotics as shown in Fig. 3E. In particular, 84 metabolites were significantly different between UFP exposed males compared to females, and 40 of these were not different in control groups (Fig. 3B). The levels of 47 metabolites were higher in UFP-exposed females compared to UFP-exposed males, and 37 metabolites were more abundant in male than in female mice. Metabolites which did not show sex differences in unexposed control groups mainly included amino acids in tryptophan metabolism, lysophospholipids, monoacylglycerols, and hexosylceramides (Fig. 3E). Additionally, 79 metabolites, 52 lower and 27 higher in males, were significantly different between males and females in the UFP+AN exposure group. These included 17 unique metabolites and 45 metabolites that did not show differences in the control groups (Fig. 3D,E), including a variety of amino acids and lipids, vitamin B6 metabolism, and benzoate metabolism. Fewer differences were seen in AN exposure groups, showing a total of 49 significantly different metabolites, 23 of which were not overlapping with the baseline differences but were mostly common with other exposure conditions (Fig. 3C, E). We also observed 25 altered metabolites shared across all four conditions (control, UFP, AN, and UFP+AN) (Fig. 3F). These metabolites are not likely related to exposure as they also differed in the controls when comparing males versus females. However, the UFP+AN combined exposure produced unique changes in 17 metabolites, compared with 16 for UFP alone, and 4 for AN alone, suggesting that the most pronounced sex-specific differences in exposure responses occurred following UFP and combined UFP+AN exposures. Overall, these results suggest that two environmental stressors, UFP and AN, have different influences on plasma metabolome in male and female mice, and the effects seem to be largely distinct from the pre-existing sexual dimorphism in unexposed animals.

### Pathway analysis: Lipid and amino acid metabolism abnormalities

To gain in-depth insight into the metabolic pathways and biological functions that were affected in the animals after different exposure conditions, we examined the super pathway and sub-pathway annotations of significantly altered metabolites, as provided by Metabolon. Significantly altered metabolites associated with UFP, AN or combined UFP+AN exposure, together with the fold change and grouped by metabolite pathways are presented in Figure 4. Among the annotated metabolites, amino acid and lipid metabolism were the top two significant metabolic pathways, followed by xenobiotic metabolism, cofactors and vitamins, peptide, nucleotide, and energy pathways (Fig. 4A, B).

**Fig. 4:**
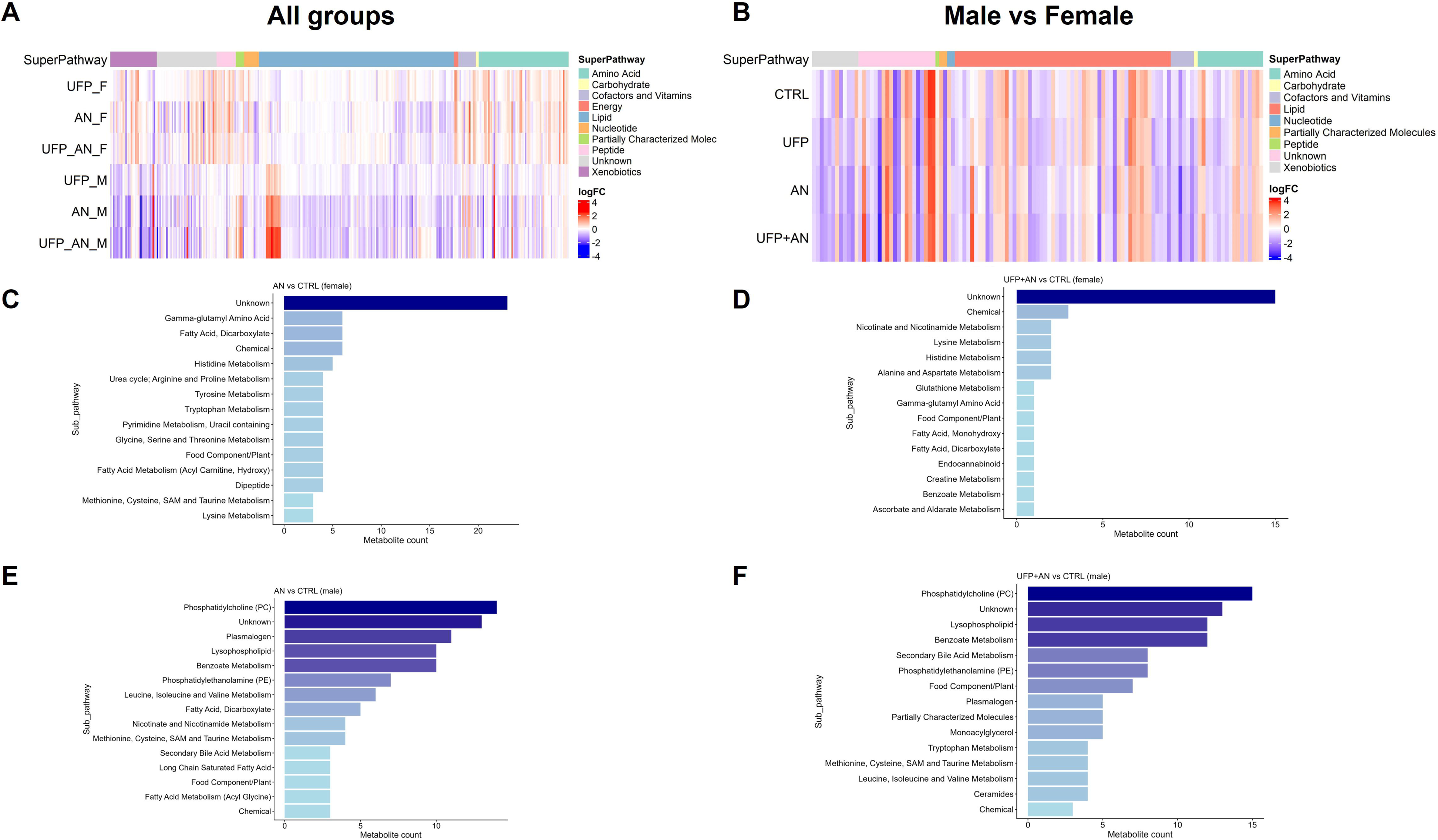
Heatmap of significantly altered metabolites across exposure groups. Log2-fold changes of A) 336 unique metabolites altered in any of the sex-stratified exposure groups, and B) 134 unique metabolites altered in any of the corresponding experimental groups between male and female mice. Metabolites are ordered based on the main biological pathways. C-F) Enriched top 15 metabolic pathways of significantly altered metabolites in females and males.

In females, the metabolomics response to exposure was primarily characterized by change in benzoate metabolism, energy metabolism, amino acid, and lipid metabolism. O-cresol-sulfate, which is involved in benzoate metabolism, was decreased in all exposure conditions (logFC –0.82 in UFP, -1.67 in AN, and –1.26 in UFP+AN exposure). Common changes were also observed, for example, in metabolites involved in TCA cycle; aconitate, malate, and citrate (logFC 0.46-1.45), and a few metabolites involved in the urea cycle (e.g., pro-hydroxy-pro; logFC 1.07-1.24) and lysine metabolism (logFC 0.36-0.66), as shown in Fig. 4C and Supplementary Table 1. Primarily in AN exposure, several gamma-glutamyl amino acids were slightly increased (logFC 0.29-0.56), while dicarboxylate fatty acids and acylcarnitines were decreased (logFC –0.26-1.07). We also observed changes in multiple amino acid pathways, such as histidine, tryptophan, and tyrosine metabolism, purine and pyrimidine metabolism (e.g, adenosine 5’-monophosphate; logFC –1.29), and an increase in dipeptides (e.g., glycylproline; logFC 1.48) in females exposed to AN (Fig. 4C). Additionally, highly altered metabolites included bilirubin (logFC 2.05) and guanidinoacetate (logFC=1.57, creatine metabolism) in AN exposure; cysteinylglycine disulfide (logFC 1.41, glutathione metabolism) in combined exposure; and N-acetyl-isoputreanine (logFC=1.39/1.14, polyamine metabolism) and naringenin 7-glucuronide (logFC –1.30/-1.47, food component/plant) in AN and combined exposures, respectively (Fig. 4D).

In males, the most affected metabolic pathways included phosphatidylcholines (PC), lysophospholipids, plasmalogen, phosphatidylethanolamines (PE), and benzoate metabolism in both AN and combined UFP+AN exposure conditions (Fig. 4E,F). The metabolites involved in these lipid and xenobiotic pathways were predominantly decreased. We also observed alterations in branched-chain amino acid metabolism, fatty acid metabolism, and nicotinate and nicotinamide metabolism, which were more prominent in AN exposure group. Conversely, highly increased secondary bile acid metabolism (logFC 1.99-2.61) and decreased food components/plant (logFC -1.10-1.69) were more extensively seen in the combined UFP+AN exposure. The most increased metabolites both in AN and UFP+AN exposure included p-cresol sulfate (logFC 3.24/2.40, benzoate metabolism), p-cresol glucuronide (logFC 3.17/2.43, tyrosine metabolism), and a secondary bile acid, 3-dehydrocholate (logFC 2.06/2.61). The most decreased metabolite was 3-(3-hydroxyphenyl)propionate sulfate (logFC –2.76/-2.49, benzoate metabolism) in both exposure conditions.

Exposure-specific sex comparisons provided further insights into sex-dependent responses. In unexposed groups, sex differences were relatively limited and mainly converged in histidine metabolism, certain fatty acids (e.g., citronellic acid, branched chain 14:0 dicarboxylic acid, and isocaproylglycine), and a few lysophospholipids, PCs and PEs along with other single metabolites. In contrast, UFP exposure expanded the sex-dependent metabolic response. In UFP exposed mice, additional sex differences were observed particularly in tryptophan metabolism and lysophospholipids, as well as in other single amino acids (e.g., phenol glucuronide), lipids (e.g., hexosylceramides, monoacylglycerols), vitamin B6 metabolism (pyridoxal), and nucleotides (guanosine). AN exposed mice showed sex differences especially in lysine metabolism, benzoate metabolism, and sphingomyelins. As expected, the same pathways differed also in the combined UFP+AN exposure.

Importantly, some metabolites showed sex differences only after exposure, suggesting the metabolic susceptibility may vary depending on sex. Notably, lyxonate, which is a carbohydrate involved in pentose metabolism, and trimethylamine N-oxide (TMAO), involved in phospholipid metabolism, showed significantly lower abundances in males compared to females after single and combined exposures, although this metabolite did not show a difference in the unexposed control groups. Furthermore, the number of significantly different cofactors and vitamins was greater under combined exposure than under single exposures or in controls. Specifically, oxalate (logFC -2.22), trigonelline/N-methylnicotinate (logFC -1.07), and pyridoxamine (logFC -0.82) were lower in males and observed only in the combined UFP+AN exposure condition.

## Discussion

In this study, we demonstrate sex-dependent systemic metabolome disruptions induced by UFP, AN and combined UFP+AN in the plasma of healthy mice, addressing an important knowledge gap regarding how air and noise pollutants differentially affect metabolic responses in males and females (Fig. 5). In comparing the different exposure groups, the data demonstrated that exposure to AN led to the largest change (measured as number of metabolites altered) in the plasma metabolome in both female and male mice. In contrast, exposure to UFP only triggered metabolite changes in female mice, highlighting a notable sex difference in the biological response to UFP exposure. Interestingly, the combined UFP+AN exposure resulted in greater number of significantly altered plasma metabolites in males than either single exposure, whereas in females the combined exposure produced a non-additive and attenuated responses compared with the individual exposures. Moreover, lipid and amino acid metabolism were identified as the top two super pathways significantly affected by exposure to UFP, AN and UFP+AN. Our analysis thereby provided evidence of sex-associated metabolic differences, which are strongest following UFP and UFP+AN exposure.

**Fig. 5:**
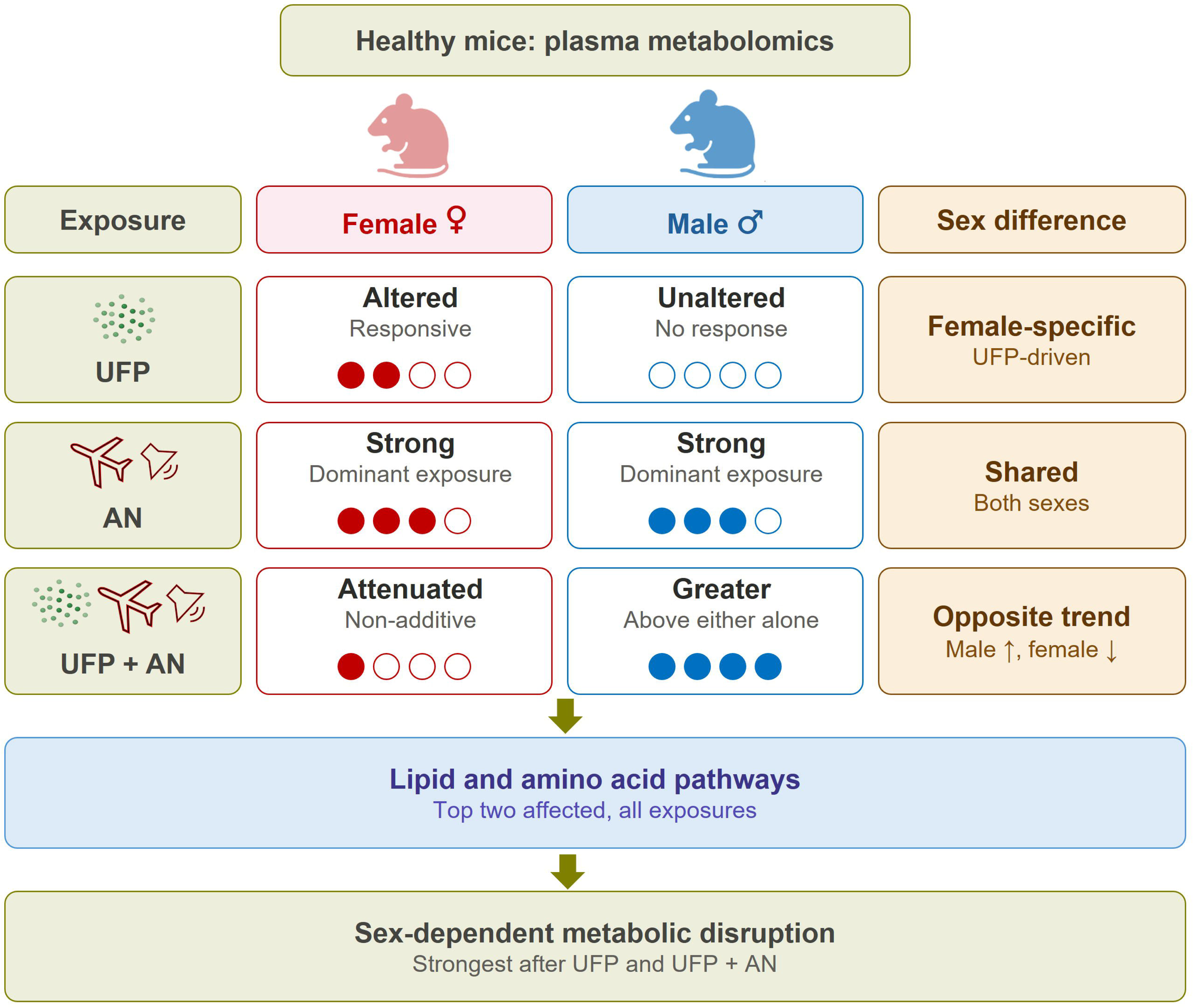
Schematic summary of sex-specific effects of UFP, AN, and UFP+AN exposure on plasma metabolic pathways.

### UFP exposure elicits female-specific metabolic responses

In real-world scenarios, individuals are often exposed to complex and simultaneous interactions among multiple hazardous factors, and the combination of different environmental exposures could therefore pose a heightened risk to health. To simulate these realistic complex exposure scenarios, we established a mouse exposure model with 9 days of exposure to UFP inhalation and AN over a period of 4 hours each day, which was informed by our prior investigations into UFP exposure ^19,20^. UFP are abundant in the atmosphere due to a combination of natural sources and processes, and those resulting from human activities ^9^. These particles can enter the human body through the olfactory system via nasal inhalation or through the lung, allowing them to translocate into different organs. Research has shown that various types of UFP can elicit similar health effects ^8,9^. For example, exposure to near-highway UFP induced various metabolic changes, including increased lipid peroxidation, which is implicated as a pro-inflammatory signaling mediators ^26^. Other epidemiological studies have also demonstrated that short-term exposure to UFP is associated with significant increases in blood pressure and alterations in metabolomic profiles in plasma and urine ^26,45^. These changes are mainly involved in systemic inflammation, oxidative stress and endothelial dysfunction. In the current study, the specific type of UFP utilized was a carbon-core UFP with an average diameter <40 nm, generated by a propane-fueled CAST-burner. Our data demonstrated that exposure to UFP alone resulted in notable shifts in plasma metabolites in females, but not males. Female plasma metabolism was specifically altered regarding pathways related to amino acids, energy metabolism, and peptide signaling. Similarly, a previous study reported that obese female mice exhibited more significant changes in both gut microbiome metabolism and plasma metabolites compared to obese male mice following gastrointestinal UFP exposure ^29^. Additionally, key pathways affected by UFP exposure were amino acids and energy metabolism ^29^. Moreover, a human cohort study of 59 adults found that long term UFP exposure was linked to altered plasma metabolomic profiles, including changes in arginine, glutamine and cystine ^26^.

### Aircraft noise and combined UFP+AN exposure

In addition to UFP exposure, environmental noise pollution presents another significant health risk, primarily affecting the body through classic stress-axis pathways ^46^. We utilized a well-established paradigm of AN, previously extensively studied by our co-authors ^21,35^. The AN paradigm used consisted of continuous repetition of a 2-hour aircraft noise recording (69 events, mean 72 dB, peak 85dB), representing a moderate exposure level below the threshold for hearing loss and allowing investigation of non-auditory health effects of AN. Earlier studies have demonstrated short-term AN (average 72dB) exposure induced functional changes in mice, such as oxidative stress, inflammation, and mitochondrial and cardiac metabolism ^35,47,48^. Recent research reported that acute exposure to high-decibel noise (120dB) severely disrupts the homeostasis of lipid metabolism in serum of rats in a time-dependent manner ^49^. In line with these studies, we found that exposure to AN at an average of 72dB for 4 h each day over 9 days caused substantial alterations in the plasma metabolome in both female and male mice. Although male and female mice shared broadly similar metabolic pathway responses to noise exposure, with only 49 metabolites differing directly between sexes, pathway enrichment revealed sex-specific patterns. Specifically, females showed greater involvement of amino acid, fatty acid, and histidine-related metabolism, whereas males showed stronger enrichment of phospholipid, plasmalogen, lysophospholipid, and benzoate-associated pathways. Consistent with this observation, an earlier study found that plasma concentrations of several unsaturated fatty acids were reduced following around-the-clock aircraft noise exposure for 4 days in male mice, indicating disruption of lipid homeostasis ^50^. Notably, in the combination of UFP+AN exposure, male mice exhibited 12.2% more altered metabolites than AN exposure alone, indicating a modestly stronger metabolomic response. The most pronounced effects were observed in lipid super pathways, where 85 altered lipids were identified in UFP+AN exposure, compared to 76 in single AN exposure. Interestingly, many affected pathways were similar in both conditions. For instance, PC, lysophospholipid, benzoate metabolism, secondary bile acid metabolisms were commonly enriched in both AN and combined UFP+AN exposure. However, combined exposure resulted in significant changes in dipeptide and bacterial/fungal-related xenobiotic pathways, which were not observed in AN exposure alone. In contrast to males, a more attenuated effect with only 45 metabolites changing was observed in female mice following UFP+AN exposure, which is substantially lower than exposure to AN alone.

### Baseline sex differences and female susceptibility to air pollution

Biological sex is one important factor influencing the human plasma metabolome, with generally modest differences in lipid, amino acid, and energy metabolism pathways, reflecting the combined effects of genetics, hormones, physiology, and lifestyle ^51^. There are also well documented differences between sexes at endocrine, physiological and behavioral levels, mainly due to sex hormone, HPA axis regulation, and body composition ^52^. While some studies on rodents have shown sex differences in plasma metabolome under normal conditions ^53–55^, there are no studies yet that explore how male and female responses to environmental stressors differ, especially regarding UFP and AN. One study highlighted differences in the plasma metabolome of unexposed adult male and female Wistar rats, including variations in amino acid, biogenic compounds, acylcarnitine ^53^. Research by Zhang *et al.* showed that 56% of the identified plasma metabolites were significantly different between the sexes in C57BL/6N mice, with most complex lipids showing higher levels in male than in female mice ^54^. In particular, 42% of measured metabolites were higher in males, including triacylglycerols, PCs and phosphatidylinositols^54^. In contrast, females had approximately 14% more of certain metabolites, such as acylcarnitines, amino acids, some bile acids and TMAO ^54^. Our study showed that 4.3% of the total detected metabolites (49 metabolites) differ by sex in 3.5 months old male and female mice. These altered metabolites are mainly lipids and amino acids. Following UFP and UFP+AN exposure, the number of significantly different metabolites increased to 7.3% and 6.9%, respectively. Approximately 50% of the metabolites did not overlap with baseline differences in the unexposed control group, which is explained by the specific pathway and response involved in UFP or AN exposure. Our data showed that the sex-difference response is driven mainly by UFP exposure, with females being more responsive. Males and females differ in body size, breathing rates, and airway dimensions ^56,57^, which might affect the deposition of UFP in the lungs or olfactory bulbs, as well as their concentration relative to body weight. Additionally, particles can also be deposited on the animal fur and thereby swallowed and ingested during grooming ^58^, introducing another layer of potential difference in response. However, some studies in humans have demonstrated that females are more vulnerable to air pollution exposure than males. A previous study found that exposure to carbon monoxide (CO), nitric oxide (NO), and nitrogen oxides (NOx) was linked to a significantly accelerated decline in bone mineral density in women, indicating their increased vulnerability to the health impacts of air pollution ^59^. Furthermore, women experienced a slightly higher risk of nonaccidental mortality and cardiovascular mortality than men following PM_2.5_exposure ^60^ ^60,61^. A recent study also showed that air pollution was significantly associated with increased cardiometabolic multimorbidity risk in women compared to men ^62^. Furthermore, a global assessment of ambient air pollution and cancer incidence reported that women exhibited consistently higher attributable risks and burdens for various cancers associated with long-term exposure to PM_2.5_, O□, and NO□ ^63^. Consistent with previous evidence, the current plasma metabolomics data support that females are more susceptible to air pollutants than males.

### Lipid metabolism

Lipids are major structural components of biological membranes and function as substrates, intermediates, and products in numerous metabolic reactions ^64^. Their metabolism is closely associated with oxidative stress, inflammation, and other biological processes. Environmental stressors, including pollutants and climate-related exposures have been reported to alter lipid profiles ^45,65–67^. For instance, a study found that exposure to road noise (over 55 dB) was associated with increased cholesterol, esterified cholesterol-rich low-density and intermediate-density lipoproteins, and specific fatty acids in the serum/plasma of three European population cohorts ^65,67^. A recent study also found that fatty acids and amino acids were significantly altered in relation to exposure to air pollution on large sample populations ^68^. Additionally, glycerophospholipid metabolism, citrate cycle, and amino acid-related pathways are associated with lung cancer incidence ^68^. Furthermore, exposure to PM_2.5_ triggered changes in plasma lipidome in a mouse model ^65^. In our study, lipid metabolism showed the most pronounced changes following UFP, AN and UFP+AN exposure in both male and female mice. Importantly, exposure to UFP, AN, or UFP+AN amplified metabolic sexual dimorphism. While baseline differences were relatively restricted in unexposed mice, UFP exposure revealed additional differences in tryptophan metabolism and lysophospholipids. AN exposure highlighted changes in lysine, benzoate, and sphingomyelin metabolism, and combined UFP+AN exposure reproduced several of these sex-dependent signatures. In males, AN and UFP+AN exposure induced predominant decreases in phosphatidylcholines, phosphatidylethanolamines, lysophospholipids, and plasmalogens. These changes suggest disruptions of membrane lipid composition, lipid remodeling, and possibly oxidative lipid damage. It is known that phosphatidylcholines and phosphatidylethanolamines are key structural components of cellular and mitochondrial membranes ^63^, while plasmalogens are phospholipids with antioxidant properties that are particularly vulnerable to oxidative stress ^69,70^. Thus, alterations in these lipids may therefore reflect compromised membrane integrity, altered inflammatory signaling, and mitochondrial dysfunctions, which are implicated in cardiovascular and metabolic diseases associated with air pollution or noise exposure ^10,18,71^. In females, AN exposure induced a broad metabolic disturbance, characterized by increased TCA-cycle intermediates, including citrate, aconitate, and malate together with changes in amino acid, urea-cycle related metabolites, lysine metabolism, and decreased acylcarnitines. A similar disturbance in acylcarnitine metabolism has been linked to mitochondrial function, insulin resistance, metabolic syndrome, and cardiovascular risk ^72^, supporting the possibility that noise exposure induces lipid metabolic changes, which may contribute to cardiometabolic vulnerability.

### Amino acid and microbiome-associated metabolism

Beyond lipid metabolism, amino acid alteration was the second most affected pathway following UFP or/and AN exposure, a finding supported by human data ^67,73,74^. Alterations in histidine/histamine, tryptophan, tyrosine, lysine, and branched-chain amino acid suggest broader disruption of neurotransmitter precursor availability, immune signaling, and host-microbiome interactions. Tryptophan metabolism is important because it connects serotonin synthesis, kynurenine pathway activation, immune regulation, and gut microbial metabolism ^75^. Within tryptophan metabolism, we observed sex differences in indoleacrylate levels in unexposed mice, suggesting inherent variations in tryptophan-associated microbial or host metabolism. Moreover, the lower levels of 5-hydroxyindoleacetate in UFP-exposed males compared to females may indicate altered serotonin turnover or stress-related neurochemical responses. Interestingly, indolelactate displayed exposure- and sex-dependent alterations. In particular, it increased in AN-exposed females, remained unchanged in females exposed to UFP+AN, and decreased in males exposed to UFP+AN. A previous study also reported that indolelactate (indolelactic acid) was significantly decreased in serum following noise-exposed male Wistar rats ^76^. Because indolelactate did not differ between sexes at baseline, our findings suggest that the observed sex differences are driven by differential responses to UFP and AN exposure rather than pre-existing baseline differences. Furthermore, the strong increases in p-cresol sulfate and p-cresol glucuronide in males, together with changes in o-cresol-sulfate in females, suggest that exposure may affect gut microbiome-derived amino acid metabolism and host phase II conjugation ^77^. Overall, our findings support a mechanistic model in which UFP, AN, and combined UFP+AN exposure may increase disease risk through interconnected disturbances in lipid and amino acid metabolism, mitochondrial functions, redox homeostasis, and microbiome-associated metabolism.

### Localized versus systemic responses to co-exposure

It should be noted that most existing studies have concentrated on lung or cardiovascular effects following co-exposure to air and noise pollution, yet most of these studies have focused on tissues, rather than systemic responses to exposure. For instance, an earlier study showed that combined exposure of PM and AN induced partially synergistically enhanced toxicity in the pulmonary, cardiovascular, and cerebral systems of male mice ^21^. Furthermore, research focusing on other co-exposure models ^76^ ^78,79^, such as ozone (O□) and PM2.5, has proposed that simultaneous exposure to adverse environmental factors may exacerbate allergic airways disease in female mice ^79^. Additionally, a recent study employing a co-exposure model of UFP with a size of approximately 30-40nm and microorganisms reported synergistic toxicological effects in the lung of male C57BL/6 mice that exceed those of either agent alone ^80^. Moreover, in the downstream analysis from lung, heart, brain tissue from animals in Kuntic *et al.* study ^21^, the authors observed constrained effects in the proteomic dataset of lung tissue of male C57BL6/J following PM+AN exposure ^22^. Ameli *et al.* reported that proteins related to ribosome/translation, RNA-processing, and chromatin-associated categories were constrained in PM+AN exposure. This effect may be attributed to a ceiling effect on biosynthetic modules when one stressor has already activated these modules, combined exposure does not result in higher protein expression ^22^. Altogether, these findings underline the complexity of UFP+AN interacting at the metabolic and molecular levels, as well as the importance of considering both localized and systemic responses in future research.

### Limitations and future directions

Although the present study primarily focused on the interplay of UFP and AN co-exposure on the systemic metabolomic response, this study has some limitations. First, we exposed mice to a relatively low dose of freshly generated UFP (mass concentration 73.2 µg/m³) for 9 days over a 2-week period and subsequently sacrificed them on the day following the final exposure day. This timeline might not capture some of the acute stress responses resulting from exposure. Additionally, we were not able to analyze the hormonal changes in our animals during exposure, which complicates the interpretation of metabolomics results as hormonal shifts can play a significant role in the sex-dependent response to environmental stressors. Furthermore, the UFP we used in the exposure mainly consists of black carbon along with a small proportion of organic compounds. This composition may differ significantly from the real-world exposure to air pollution, which often contains a vast variety of different compounds, metals and other toxic substances known to trigger health risks ^4,9,81^. Future studies need to analyze the hormonal response and extend the exposure periods to gain comprehensive understanding of the chronic effects of UFP on metabolism. Additionally, further analysis of specific tissues or organs in both sexes to identify the molecular and cellular mechanisms in different biological systems under exposure will enhance our understanding of the health implications associated with air and noise pollution in real-world scenarios.

## Conclusion

In conclusion, this study highlights that exposure to UFP, AN, and combined UFP+AN induces significant systemic metabolomic changes in healthy mice, with marked sex-dependent differences. Lipid and amino acid metabolism are the most affected pathways, indicating disruptions in membrane lipid homeostasis, mitochondrial energy metabolism, fatty acid handling, and amino acid turnover. These findings provide evidence that the interplay between air pollution particles and aircraft noise can induce systemic metabolic alterations in otherwise healthy animals, potentially linked to oxidative stress, inflammation, mitochondrial dysfunction, and risks of cardiometabolic or neuroendocrine diseases.

## Credit authorship contribution statement

**Thuy Thi Lai**: Writing-original draft, writing-review and editing, conceptualization, data curation, formal analysis, investigation, methodology, project administration, validation, visualization. **Aino-Kaisa Piironen**: Writing-original draft, writing-review and editing, conceptualization, data curation, formal analysis, validation, visualization, **Elena Puris:** writing-review and editing, conceptualization, validation. **Veronika Górová**: writing-review and editing, investigation. **Tuukka Kokkola**: writing-review and editing, investigation, formal analysis. **Mika Ihalainen**: writing-review and editing, investigation. **Jiayin Zheng**: writing-review and editing, methodology. **Thomas Münzel**: writing-review and editing, methodology. **Andreas Daiber**: writing-review and editing, methodology. **Pasi Jalava**: writing-review and editing, methodology, conceptualization. **Olli Sippula:** writing-review and editing, methodology, conceptualization. **Katja M. Kanninen:** Writing-original draft, writing-review and editing, supervision, resources, methodology, conceptualization, funding acquisition.

## Declaration of generative AI use

During the preparation of this work, the authors used Copilot and Grammarly to assist with language editing and improve readability. After using these tools, the authors reviewed and edited the content as needed and take full responsibility for the content of the publication. No generative AI or AI-assisted tools were used to generate data, perform analyses, create figures, or draw scientific conclusions.

## Data availability

All data and code will be deposited in a public repository (e.g., GitLab, GitHub,) upon publication.

## Acknowledgements

The authors would like to thank Laura Mussalo, Hennariikka Koivisto, Pasi Miettinen, Mirka Tikkanen, Luca Soraru and animal caretakers at the Lab Animal Center for their assistance.

