## Supplementary Fig. 1 for "Plasma untargeted metabolomics reveals distinct metabolic disturbances between sexes following inhalation of ultrafine particles and exposure to aircraft noise in mice"

**Supplementary Figure**

**Supplementary Fig. 1: Experimental design and exposure characterization for UFP and AN** A) Schematic illustration of the setups for particulate matter generation and animal inhalation chamber. B) TEM image of particles. C) Particle size distribution. D) Aircraft noise exposure pattern. E-G) Animal weights at baseline and after 1- exposure and endpoint.

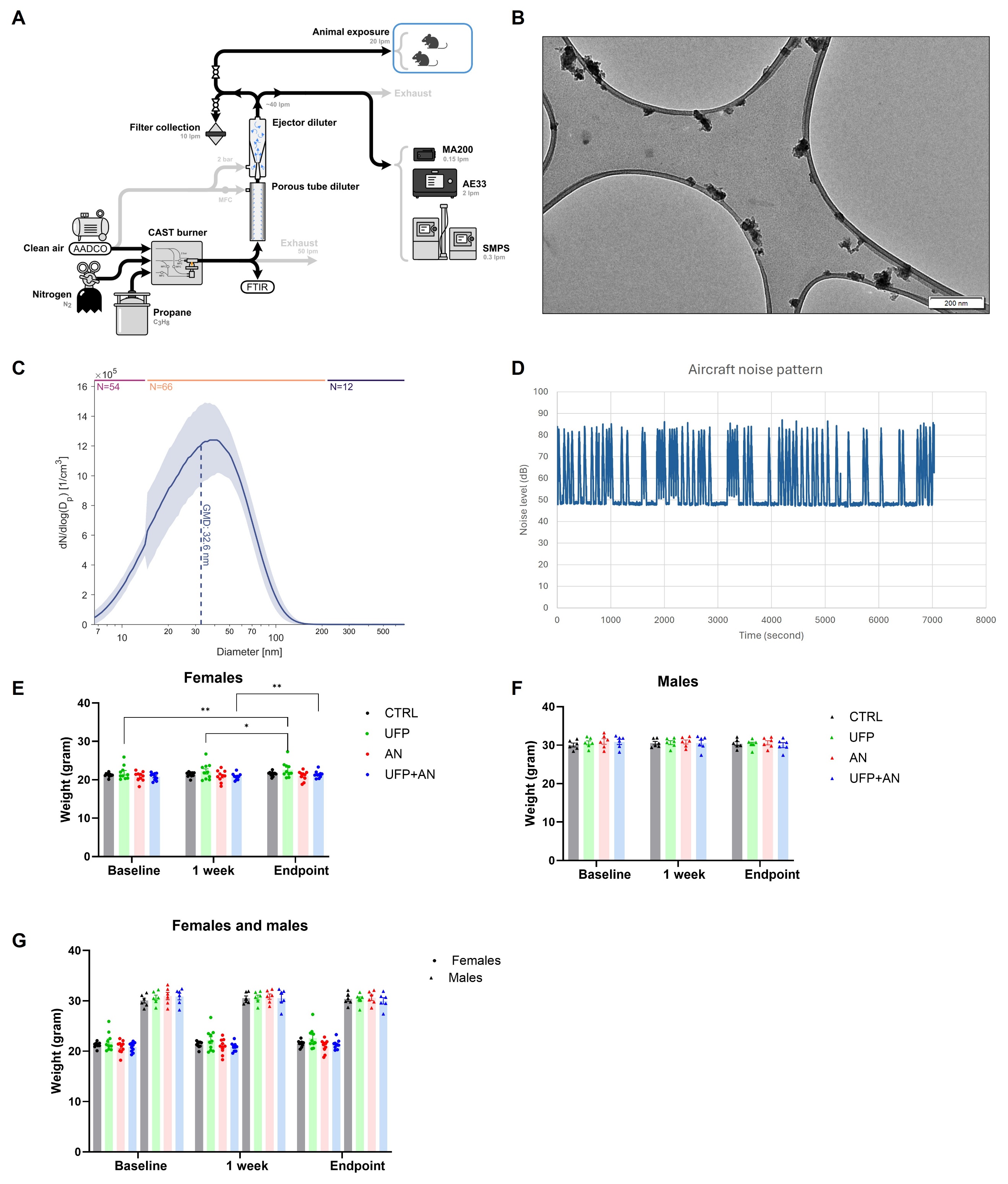
